# Improving power for rare variant tests by integrating external controls

**DOI:** 10.1101/081711

**Authors:** Seunggeun Lee, Sehee Kim, Christian Fuchsberger

## Abstract

Due to the drop in sequencing cost, the number of sequenced genomes is increasing rapidly. To improve power of rare variant tests, these sequenced samples could be used as external control samples in addition to control samples from the study itself. However, when using external controls, possible batch effects due to the use of different sequencing platforms or genotype calling pipelines can dramatically increase type I error rates. To address this, we propose novel summary statistics-based single and gene- or region-based rare-variant tests that allow the integration of external controls while controlling for type I error. Our approach is based on the insight that batch effects on a given variant can be assessed by comparing odds ratio estimates using internal controls only vs. using combined control samples of internal and external controls. From simulation experiments and the analysis of data from age related macular degeneration and type 2 diabetes studies, we demonstrate that our method can substantially improve power while controlling for type I error rate.

## Introduction

Identification of genetic variants that predispose to complex diseases is an essential step toward understanding disease etiology, which can lead to breakthroughs in diagnosis, prevention, and treatment. The advance of sequencing technologies (Shendure & Ji, 2008) enables studying the full spectrum of genetic variants, including rare variants (minor allele frequency (MAF) < 1%), in large studies. Although recent sequencing studies have started to identify disease-associated rare variants (Cruchaga et al., 2014; Steinthorsdottir et al., 2014), the number of discoveries is much smaller than initially predicted (Zuk et al., 2014). To facilitate further discoveries, more efficient and powerful statistical strategies and methods are needed.

Given decreasing sequencing costs, the number of sequenced whole exomes and even whole genomes are rapidly increasing. These sequenced samples provide a great opportunity to increase the power of rare variant test. For a single study, if sequenced samples from other studies are used as control samples (external control samples), the power of rare variant tests can be substantially improved without any additional sequencing cost. For example, using 1,529 external control samples from NHLBI ESP data in addition to their own 789 internal control samples, Zhan et al. (2013) identified a rare coding variant (MAF=0.4%, p-value=2.7×10^−4^) in the C3 gene associated with age-related macular degeneration. When they exclusively used internal controls, the variant was substantially less significant.

Although the use of external control samples can greatly improve power, systematic differences between studies can have a negative impact on type I error control and power. Study heterogeneity can arise from differences in study populations, i.e. population stratification, or technical batch effects due to the use of different sequencing platforms or genotype calling pipelines (Quail et al., 2012). Recently, substantial progress has been made in identifying external control samples with similar genetic background (Wang et al., 2014); however, only limited success has been made in adjusting for technical batch effects. Derkach et al. (2014) developed a robust score test that uses genotype likelihoods. The method can control type I error rates when locations of variants are correctly known; however, inflated type I error rates are noted when variant locations should be inferred from genotype calls (Hu, Liao, Johnston, Allen, & Satten, 2015). More recently, Hu et al. (2015) developed a likelihood-based burden test that uses sequence reads, which can provide more accurate type I error rate controls. However, given that raw high depth sequence read data can be > 1000 times larger than processed genotype data, downloading, storing, and analyzing external control sequence read data can be a huge computational burden. In addition, the aforementioned methods need individual-level genotype and phenotype information, which is often difficult to obtain.

To address these challenges, we propose simple and robust rare variant association tests that only require allele counts from external studies. Our approach, integrating External Controls into Association Test (iECAT), is based on the insight that the existence of batch effects on a given variant can be assessed by comparing two different odds ratio estimates: odds ratio estimate using internal controls only vs. odds ratio estimate using combined control samples of internal and external controls. If a variant is subject to batch effects, these two should be substantially different; even when ancestry-matched external controls were used (Mahajan & Robertson, 2015). In such a case, only internal control samples should be used to avoid false positives. Otherwise, external control samples can be added to increase the sample size. To approach this problem data-adaptively, we propose using an empirical Bayesian-type method. We first construct a single variant test based on the shrinkage method and extend it to several gene- or region-based tests, such as burden tests (Li & Leal, 2008; Madsen & Browning, 2009; A. P. Morris & Zeggini, 2010), variance-component tests (Wu et al., 2011), and combinations of these two (e.g. SKAT-O (Lee, Wu, & Lin, 2012)).

Given that only allele counts from external studies are needed for the proposed method, the method can be used with summary information publicly available in variant servers, such as the ESP Exome Variant Server. Through extensive simulation studies and analysis of AMD and type 2 diabetes sequencing data, we demonstrate that the proposed method can improve power while controlling for type I error rates.

## Material and Methods

### Single variant association test

For a single-variant test, we consider the following shrinkage estimation-based allelic test with an assumption that the Hardy-Weinberg equilibrium (HWE) holds. Let *Y*=1 (*Y*=0) denote affected (unaffected) status, *G*=1 (*G*=0) denote the minor allele (major allele) in the variant. Let 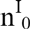 and 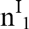 (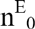 and 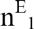) denote the number of controls and cases of an internal study (external study), respectively. The data are represented by a 3 × 2 table in Table 1.

**Table 1.** Data setup for internal case-control and external control samples for a single variant association test.

| | | $G_j=1$ | $G_j=0$ | Total |
| --- | --- | --- | --- | --- |
| Internal | $Y=1$ | $r^I_1$ | $2n^I_1 - r^I_1$ | $2n^I_1$ |
| | $Y=0$ | $r^I_0$ | $2n^I_0 - r^I_0$ | $2n^I_0$ |
| External | $Y=0$ | $r^E_0$ | $2n^E_0 - r^E_0$ | $2n^E_0$ |

Considering the internal study only, let 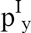 denote the *unknown true* minor allele frequency (i.e., 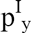 = Pr[G=1|Y=y] for y=0, 1), and 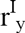 denote the corresponding *observed* number of minor alleles. The observed counts can be viewed as a random sample from two independent binomial distributions, 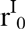 ~ Binomial(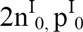) and 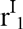 ~ Binomial(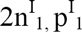). Note that the number of trials is two times 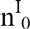 (or 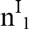) since each sample consists of two copies of chromosomes. Suppose the parameter of interest is a log odds ratio of the genetic effect, given by

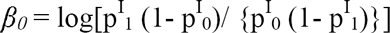

Our goal is to test whether 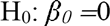 or not. By using the internal study only, 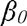 can be consistently estimated by 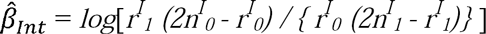.

Now, suppose 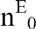 external control samples are available, and assume the *observed* number of minor alleles, 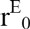, follows a binomial distribution with 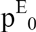, the proportion of the minor allele in the external controls, i.e., 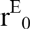 ~ Binomial(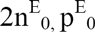). Combining control samples from internal and external studies, the estimate of 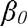 is

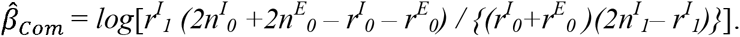

When 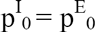, indicating that no batch effect exists, 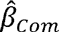 converges to 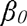 in probability. When 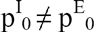, however, 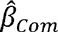 converges to a biased quantity, 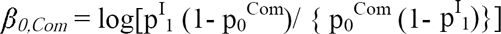, where 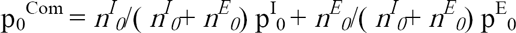. Thus, the bias can be quantified as 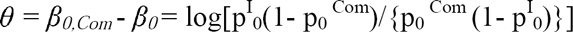. Taking it for granted that the bias would exist, we propose a bias-adjusted estimate of 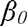 as follows.

We first estimate the bias *θ* by using an empirical Bayesian approach. It begins with an estimate 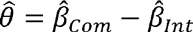 that asymptotically follows a normal distribution 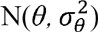, where 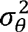 is the variance of 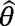. To account for the uncertainty in the bias, we assume that *θ* is a random variable following a normal distribution N(0, *v*), where the unknown hyperparameter, *v*, reflects the magnitude of uncertainty by the batch effects. Note that the consistent estimator of *v* from the marginal distribution of 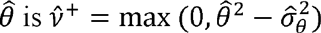, where 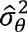 is an estimate of 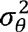 (C. N. Morris, 1983). Recently Mukherjee and Chatterjee (2008) have proposed to use 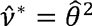, a more conservative estimator of *v*. The main advantage of using 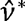 instead of 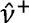 is that 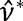 provides a closed-form of variance estimator of the subsequent *θ* estimator. They also showed that there is essentially no loss of efficiency of using 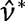. Therefore, we propose to use 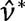, and the resulting estimate of the posterior mean of *θ* is 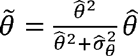. Finally, the proposed estimator of 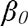 is

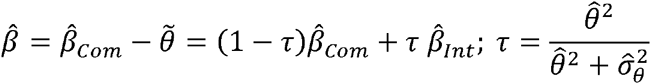

When 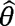 (i.e., the difference between 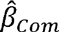 and 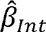) is large, *τ* is close to one, and hence 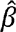 becomes 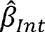. In contrast, when *τ* is close to zero due to a small 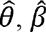 is close to 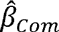. Regardless of the size of the true bias 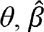 is an asymptotically unbiased estimator of 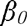 (Web Appendix A). The exact expression of the asymptotic variance of 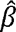 can be found in the Web Appendix A.

If there is a prior information on when *θ* could be zero under the null hypothesis (i.e 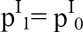), using *τ* = 0 can increase power without inflating type I error rates. Suppose that observed MAFs of external controls (i.e. 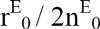) is in between observed MAFs of cases and internal controls (i.e. 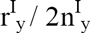), and hence 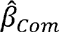 is closer to zero than 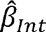 (i.e. 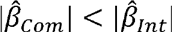). Under the null hypothesis with the batch effects (i.e. 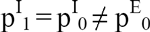), since observed MAFs converge to true MAFs, the probability to observe this phenomenon will converge to zero. This indicates that using *τ* = 0 in this situation will not increase type I error rates. In contrast, using *τ* = 0 in this situation will increase power especially when the external controls include case samples, since the case contamination will cause either 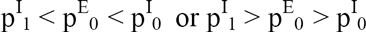 under the alternative hypothesis. And hence when this phenomenon occurs, we used *τ* = 0 as a default value in simulation studies and real data analysis.

### Gene- or region-based test

For gene- or region-based tests, we extend the proposed single-variant test in the previous section to Burden, SKAT and SKAT-O type tests. The key idea is first to construct a single-variant score-type statistic for each variant using 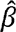 and then to aggregate them using their weighted linear or quadratic sums. To aggregate association signals in each variant, the Burden test uses a weighted linear sum of score statistics, whereas SKAT uses a weighted quadratic sum of score statistics. The combined test, SKAT-O, uses a linear combination of the Burden and SKAT test statistics (Lee, Abecasis, Boehnke, & Lin, 2014).

Suppose that a region being tested has *p* variant loci. For variant *j*, let 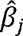 be the log odds ratio estimate in (1), 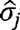 the standard error estimate, and *q*_*j*_ the MAF estimate from the internal samples. In Web Appendix B, we show that a single variant score statistic is approximately proportional to the product of log odds ratio estimate, sample size and the genotype variance. Using this fact, we construct a score-type statistic for a single variant *j* as 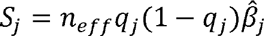, where *n*_*eff*_ is the effective sample size, and propose the following test statistics:

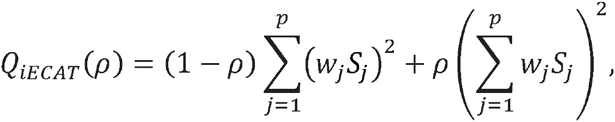

where *w_j_ is* a weight for variant *j*, and *ρ* is a parameter between 0 and 1. Clearly, *ρ=0* corresponds to the SKAT-type test, and *ρ=1* corresponds to the Burden-type tests. Note that 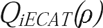 has a similar form to the variance component test with the compound symmetric kernel structure proposed by Lee et al. (2012). More details on the test and p-value calculation are given in Web Appendix C.

Burden type tests (i.e. *ρ=1*) are powerful when large percentage of variants are causal and effects are in the same direction. SKAT type tests (i.e. *ρ=0*) are powerful when heterogeneity is noted regarding effect sizes and the direction of the effects (Basu & Pan, 2011; Lee et al., 2014). SKAT-O type tests combine Burden and SKAT tests using the minimum p-values from a grid of *ρ.* We propose the following SKAT-O type combined test as

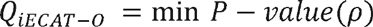

where *P-value(ρ)* is the p-value 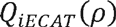 with a given *ρ.* We used a grid of eight *ρ* values *(0, 0.1^2^, 0.2^2^, 0.3^2^, 0.4^2^, 0.5^2^, 0.5, 1)* in simulation studies and real data analysis. This approach has been used in previous studies and shown to provide good performances in type I error control and power (Lee, Teslovich, Boehnke, & Lin, 2013). P-values of 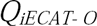 can be obtained using numerical integration, as described in Lee *et al.* (Lee et al., 2012).

### Type I error and power simulations

We performed extensive simulation studies to evaluate the performance of the proposed iECAT method. To generate sequence information, we simulated 40,000 European-like and 40,000 African-America-like haplotypes for 200 kbps using the coalescent simulator COSI with the calibrated demographic model (Schaffner et al., 2005). The binary phenotypes were generated from the following logistic regression model:

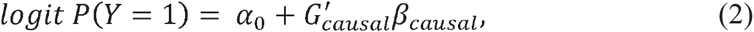

where G_causal_ is a genotype vector containing causal variants, and 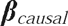 is a vector of the genetic effect coefficients. The intercept *α*_*0*_ was chosen for the disease prevalence of 0.05.

We presumed that 3% of the variants had different MAFs between internal and external control samples due to batch effects, mimicking the level of the batch effects observed in the real data analysis (see Results section for details). For these variants, MAFs in external control samples were randomly generated from Uniform (0.1×q, 4×q), where q is the MAFs in internal study samples. To mimic realistic scenarios of population stratification between internal and external control samples, we generated 0% (no population stratification), 5% and 10% of external control samples from the African-American-like haplotypes. All other internal and remaining external control samples were simulated from the European-like haplotypes.

We compared the following methods for a region-based test: 1) the proposed iECAT-O; 2) SKAT-O using the internal control only; 3) iECAT-O without adjusting for batch effects (iECAT-O_Noadj_) – that is, 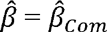 was always used regardless of the existence of the batch effects; 4) iECAT-O assuming that batch effect variants were known prior to the test (iECAT-O_Known_) – that is, 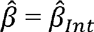 was used (i.e., *τ* is fixed with 1) in the presence of true batch effects. Otherwise, 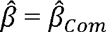 was used (i.e., *τ* is fixed with 0). Since the test iECAT-O_Known_ was constructed given that variants subject to batch effects were fully known, iECAT-O_Known_ could reach the theoretical maximum power under the iECAT framework. Obviously, this batch effect information could not be known in real data. Therefore, iECAT-O_Known_ will be used only for power comparison to quantify the efficiency loss in iECAT-O due to the uncertainty in τ. For a single variant test, we used aforementioned approaches (iECAT-O, iECAT-O_Noadj_ and iECAT-O_Known_) and the Wald test (internal controls only) with testing one variant at a time.

For single variant tests, we randomly selected a variant and tested for an association between the variant and the phenotypes. For region-based tests, we randomly selected a 3 kbp region and tested for an association between variants in the region and the phenotypes. For very rare variants, the proposed methods cannot be used because 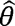 estimates can be unstable. We thus only used variants observed in both internal and external studies with internal study MAC > 3.

In type I error simulations, phenotypes were generated from (2) with 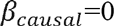. For each simulation, to evaluate type I error rates at genome-wide significance levels, we generated 10^7^ datasets for region-based tests and 5×10^7^ datasets for single variant tests. Given that generating these large datasets was computationally intensive, we considered only two internal study sample sizes, moderate and large (N_internal_=4000 and 10,000). In each sample size, two different ratios of case-control (1:1 and 2:1) were considered. The external control sample size (N_external_) was set to be the same as the internal study sample size.

For power simulations for gene-based tests, 5%, 10%, or 30% of rare variants (MAF < 1%) were assumed to be causal. In each setting, either all causal variants were risk-increasing or 80% of causal variants were risk-increasing (the remaining 20% were risk-decreasing). Given that it is possible that rarer variants have larger effect sizes, we modeled log OR as a decreasing function of MAF. Specifically, 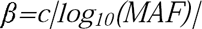. When 30% of variants were causal, we used c= log(2)/2, which led to OR=2 for a causal variant with MAF=1%. When 5% or 10% of rare variants were causal, we used c= log(4)/2 and c= log(3)/2 to compensate the decreased numbers of causal variants. For single variant tests, we evaluated power for testing a variant with MAF=(0.01, 0.005) and OR=2.

To evaluate the power when external control samples contain samples with the disease of interest (i.e., case-contamination), we generated external control samples of which 0%, 5% and 10% were the diseased samples. Because 5% disease prevalence was noted, 5% contamination was equivalent to using the general population as external controls. In all power simulation scenarios, the external control sample size was the same as the internal study sample size (N_external_=N_internal_). For each power simulation setting, we generated 5000 data sets.

### Real data analysis

We applied our method to two sequence datasets while using 4300 ESP European samples as external controls: targeted sequencing data from the age-related macular degeneration (AMD) study (Zhan et al., 2013) and deep whole exome sequence data from the Genetics of Type-2 Diabetes (GoT2D) study. The AMD study sequenced 10 target loci spanning 56 genes (2.7 Mb in total). We downloaded 3350 mapped SRA files from dbGaP (Tryka et al., 2014) (phs000684.v1.p1), applied variant calling and QC procedures as described in Zhan et al. (2013), and retained 2317 cases and 791 controls. The GoT2D study whole exome sequenced 1,326 European T2D cases and 1,331 European T2D controls at high depth. For both datasets, we applied LASER software with the HGDP reference (Wang et al., 2014) to identify population structure. Given that Finish samples were separately clustered from other European samples (Web Figure 8), and ESP has only a small number of samples clustered together with Finish samples (Web Figure 8), we exclusively used the non-Finish cohorts (British, German and Sweden) in the GoT2D data analysis, leading to a total of 650 cases and 646 controls.

For external controls, allele counts of ESP samples were downloaded from the ESP Exome Variant Sever. Variants observed in both internal and external studies were included in the analysis, after excluding variants that did not pass each study’s own QC criteria. We further excluded variants with internal study MAC ≤ 3. For gene-based tests, we used genes with at least 3 rare variants. To investigate whether ancestry matching can improve type I error control and power for whole exome scale data analysis, we obtained individual-level genotype data of ESP for the GoT2D data analysis. We used LASER software with the HGDP reference (Wang et al., 2014) for 1:2 matching between internal and external samples and identified 2,568 ancestry-matched external control samples (Web Figure 8).

## Results

### Type I error rate simulation results

The empirical type I error rates estimated for iECAT-O and the other methods are presented in Table 2 for α = 10^−4^ and 2.5×10^−6^, corresponding to candidate gene studies of 500 genes and exome-wide studies of 20,000 genes. All internal and external samples were simulated using European-like haplotypes. In all the simulation scenarios we considered, iECAT-O had well controlled or slightly inflated type I error rates. As expected, when we used the combined control samples but ignored batch effects (iECAT-O_Noadj_), type I error rates were significantly inflated. For example, empirical type I error rates were increased more than 10000–fold compared to the nominal *α* level on average when α =2.5×10^−6^. When we exclusively used the internal control samples (SKAT-O), type I error rates were well controlled.

**Table 2.** Empirical type I error rates for iECAT-O, SKAT-O and iECAT-O_Noadj_. Each cell has an empirical type I error rate estimated from 10^7^ simulated datasets. The external control sample sizes were the same as the internal sample sizes. All internal and external controls samples were simulated from European-like haplotypes.

| Internal | Internal |  |  |  |  |
| --- | --- | --- | --- | --- | --- |
| Sample Size | Case:Control | Level $\alpha$ | iECAT-O | SKAT-O | iECAT-O<sub>Noadj</sub> |
| 4000 | 1:1 | $10^{-4}$ | 7.40E-05 | 1.10E-04 | 7.00E-02 |
| | 1:1 | $2.5 \times 10^{-6}$ | 1.40E-06 | 2.10E-06 | 5.50E-02 |
| | 2:1 | $10^{-4}$ | 8.00E-05 | 1.20E-04 | 8.40E-02 |
| | 2:1 | $2.5 \times 10^{-6}$ | 2.60E-06 | 2.10E-06 | 7.00E-02 |
| 10000 | 1:1 | $10^{-4}$ | 8.60E-05 | 1.10E-04 | 1.10E-01 |
| | 1:1 | $2.5 \times 10^{-6}$ | 3.50E-06 | 2.90E-06 | 9.30E-02 |
| | 2:1 | $10^{-4}$ | 8.70E-05 | 1.10E-04 | 1.20E-01 |
| | 2:1 | $2.5 \times 10^{-6}$ | 2.40E-06 | 3.10E-06 | 1.10E-01 |

When only summary-level information for external control samples is available, samples with different genetic background cannot be excluded. We carried out additional type I error simulations with 5% and 10% of external control samples from the African-American-like haplotypes, and the results have shown that iECAT-O had robust type I error control even in the presence of population stratification (Details See Web Appendix D).

### Power simulation results

Next, we compared power of the proposed and existing methods to identify genetic associations at α= 2.5×10^−6^. Figure 1 presents power simulation results when all causal variants were deleterious variants. iECAT-O had substantially improved power compared to the approach using internal study samples only. For example, when the internal study case-control ratio was 1:1 and the internal study sample size was 10,000, iECAT-O was on average 38% to 60% more powerful. When the internal study case-control ratio was 2:1, iECAT-O exhibited increased power. On average, when the internal study sample size was 10,000, the strategy to sequence more cases (2:1 case-control ratio compared to 1:1) resulted in 11% to 17% increased power for iECAT-O compared to sequencing the same number of cases and controls (Figure 1, bottom panel). As expected, the power of iECAT-O_Known_, where the optimal value of shrinkage weight *τ* was used, was higher than that of iECAT-O, but the difference was not substantial. This implies that estimating *τ* in iECAT-O method did not reduce the efficiency so much. Given that iECAT-O_Noadj_ greatly increased type I error rates, we did not include it in this plot. The relative performance of the methods in the presence of both risk-increasing and risk-decreasing variants was quantitatively similar (Web Figure 2).

**Figure 1.**
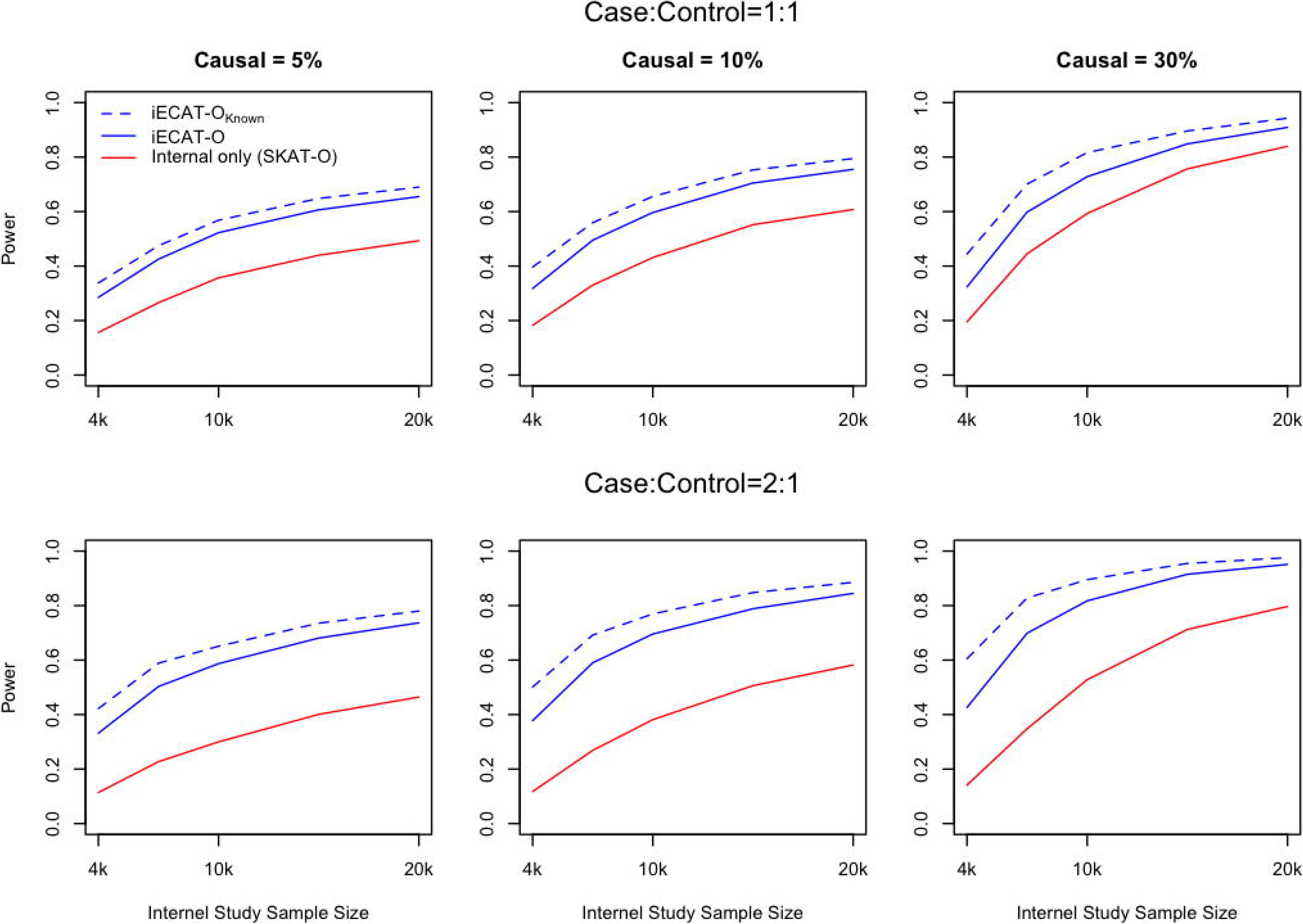
Power comparisons when all causal variants were risk increasing variants. Each line represents empirical power at α = 2.5×10^−6^. From left to right, the plots consider that 5%, 10%, and 30% of variants were causal variants, respectively. From top to bottom the plots consider that internal study case:control ratio was 1:1 and 2:1, respectively. The external control sample sizes were the same as the internal study sample sizes. For causal variants, we assumed that 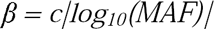 (See Method section).

**Figure 2.**
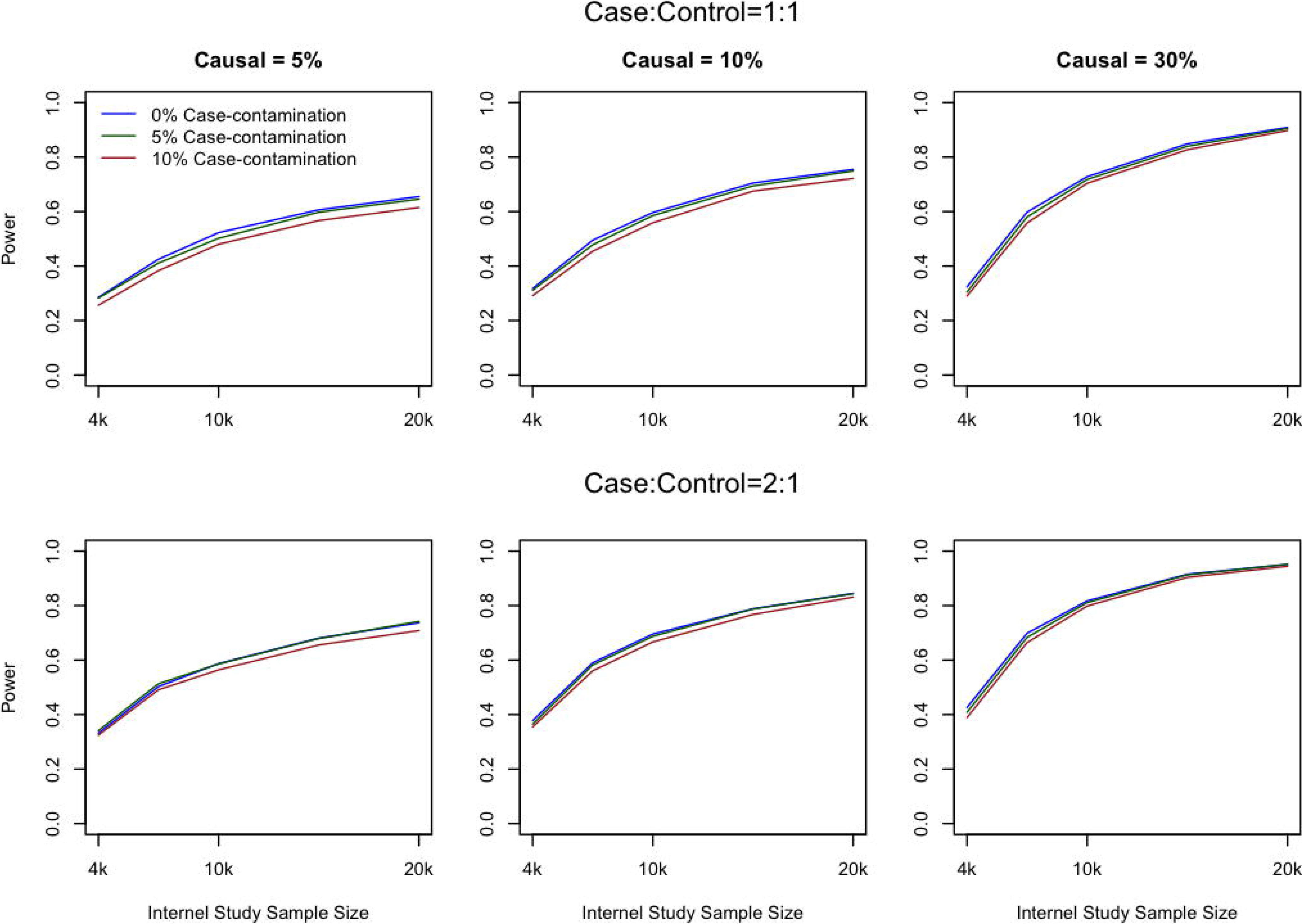
Empirical power of iECAT-O in the presence of case-contamination in external controls. Each line represents empirical power at α = 2.5×10^−6^. From left to right, the plots consider that 5%, 10%, and 30% of variants were causal variants, respectively. From top to bottom the plots consider that internal study case:control ratio was 1:1 and 2:1, respectively. All causal variants were risk-increasing variants. The external control sample sizes were the same as the internal study sample size. For causal variants, we assumed that 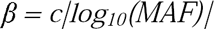 (See Method section).

Given that external studies were not performed for the target disease of interest in the internal study, it is possible that external control samples included cases (i.e., the diseased samples), and this case-contamination can reduce the power. Figure 2 presents power evaluation results with 0%, 5% and 10% of the external control samples being cases. Given that 5% disease prevalence was assumed, 5% contamination is equivalent to using the general population as external controls. When N_internal_=10,000, the power was decreased on average by 2 to 4% for 5% contamination and 3 to 8% for 10% contamination. When the sample size was small, the power reduction was slightly larger.

Next, we performed power simulations in the presence of population stratification between internal and external samples in which 5% and 10% of the external control samples were generated from African American-like haplotypes and all other samples were generated from European-like haplotypes. Given that population stratification can inflate type I error rates, we used empirical α levels obtained in the type I error simulation studies for fair comparisons. Web Figure 3 indicates that the power of iECAT-O slightly decreased in the presence of population stratification. These results suggest that the power of iECAT-O can be improved if the ancestry-based matching method is used to exclude external control samples with different genetic backgrounds.

**Figure 3.**
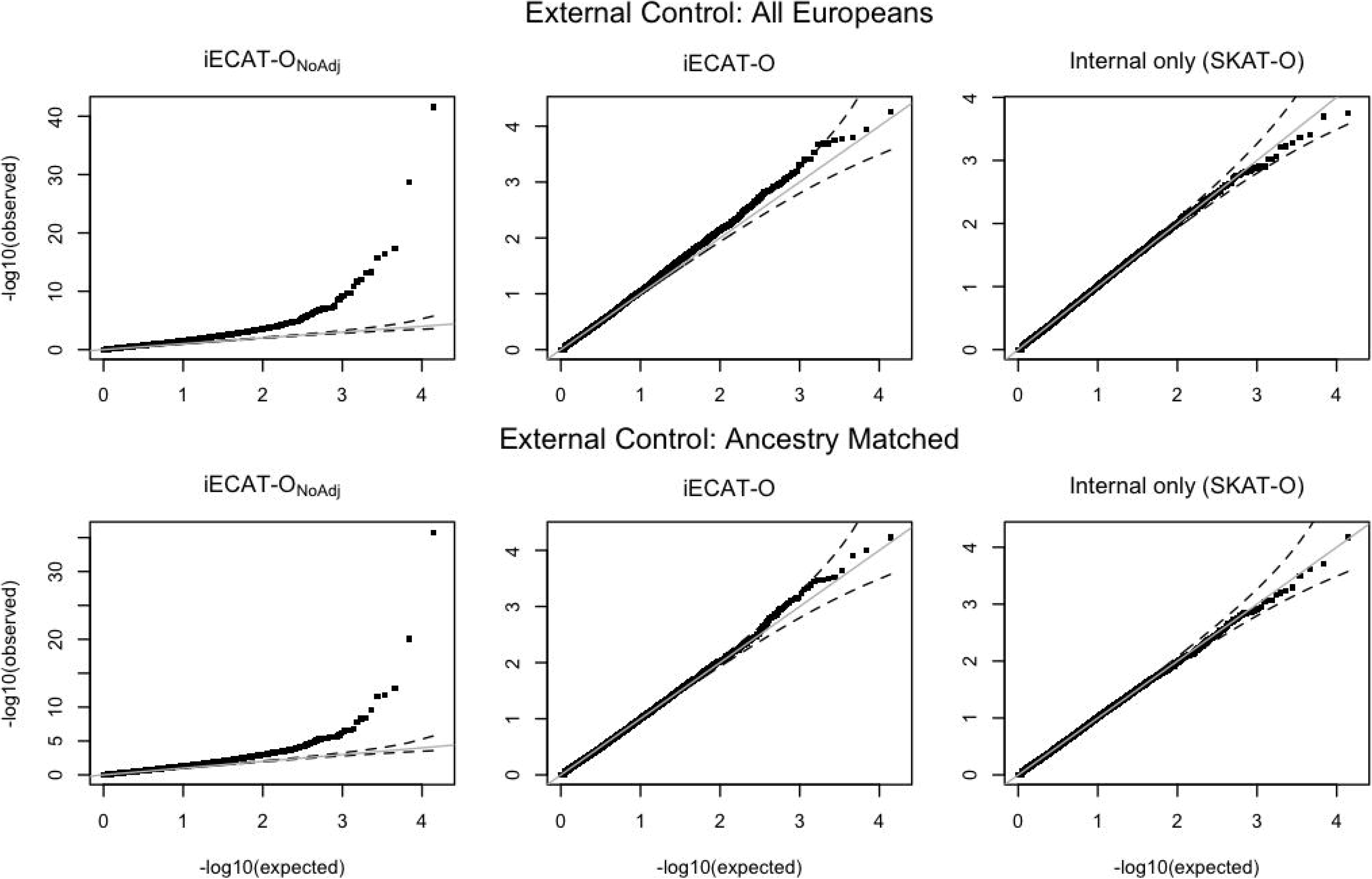
Analysis of the GoT2D exome data with ESP as external control samples. QQ-plots of -log10 p-values of gene-based tests for rare variants (MAF < 0.01). A total of 11283 genes with at least three rare variants were tested for associations with T2D status. The dashed line represents a 95% confidence band. Top panel: all 4300 European ESP samples were used as external controls. Bottom panel: 2568 ancestry matched ESP samples were used as external controls.

As expected, the power simulation results with for iECAT with *ρ*=0 (SKAT type test) and *ρ*=1 (Burden type of tests) were quantitatively similar (Web Figure 4-5). We obtained power with empirical α levels instead of nominal α levels for N_internal_ =4000 and 10,000, and the results were also quantitatively similar (Web Figure 6). We then performed power simulations for single variant tests for two different MAFs (0.01 and 0.005) (Web Figure 7). The power curves revealed that iECAT had greatly improved power compared with the method exclusively using internal controls (e.g., 43% vs. 12% when MAF=0.5%, N_internal_=10,000 and case:control=1:1). Interestingly, the strategy of sequencing more cases (case:control=2:1) did not always improve the power compared with the one case per one control strategy. For example, when MAF=0.005 and N_internal_ =15,000, case:control=2:1 design had 72% power, while case:control=1:1 design had 75% power. This occurred in part because having a smaller number of internal controls increased the variability of the bias estimate, which had a negative impart on power. We note that this increased variability had a smaller effect on gene-based tests given that the gene-based approach combines multiple single variant odds ratios.

Overall, our simulation studies demonstrate that iECAT can substantially improve power compared with the existing approaches that exclusively use internal control samples, while controlling for type I error in a wide range of scenarios. In contrast, if the external control samples are used without adjusting for batch effects, it can result in significantly increased type I error rates.

### Real data analysis

We applied our proposed methods to the analysis of AMD targeted sequencing and GoT2D whole exome sequencing datasets. For both datasets, we used allele count statistics of 4300 ESP European samples downloaded from Exome Browser Server as external controls. In addition, for the GoT2D data analysis we obtained the ancestry-matched external control samples using individual-level genotype data of ESP to investigate whether matching can improve type I error control and power in whole exome scale analysis.

#### AMD Study

We applied iECAT to 56 candidate genes in the AMD targeted sequencing dataset. After quality control, we retained 2317 cases and 791 controls. Table 3 presents the top 5 genes by iECAT-O. For this analysis, we focused on rare variants (MAF < 0.01). By combining the internal and external controls, iECAT-O revealed that two well-known AMD-related genes, *C3* and *CFH*, had the smallest p-values (p-value=5.75×10^−6^ and p-value=7.44×10^−6^, respectively), and the p-values were still significant after the Bonferroni correction (corrected α= 0.05/56=8.9×10^−4^). In contrast, when SKAT-O was performed with the AMD dataset alone, the p-values were greater than 0.01, indicating that the proposed test significantly improved power. We also performed single variant tests for a total of 538 rare variants (Web Table 3). We found that SNV rs147859257 in *C3* (p-value = 1.23×10^−5^) and rs121913059 in *CFH* (p-value=1.24×10^−5^) were the top two variants by iECAT. However, when the AMD dataset was used alone, these two SNVs resulted in p-values > 10^−2^. The single variant test results were largely consistent with Zhan et al. (2013), in which 1529 ancestry-matched external controls were used.

**Table 3.** Top five genes by iECAT-O p-values from the AMD data analysis. The 4300 European ESP samples were used as external controls. Rare variants (MAF < 0.01) were exclusively used for this analysis.

| Gene | Chr | # of variant* | iECAT-O<br>P-values | SKAT-O<br>P-values |
| --- | --- | --- | --- | --- |
| CFH | 1 | 14 | 5.75E-06 | 6.70E-02 |
| C3 | 19 | 27 | 7.44E-06 | 1.04E-02 |
| DOM3Z | 6 | 7 | 1.16E-04 | 1.69E-03 |
| CFHR4 | 1 | 6 | 8.03E-04 | 6.66E-02 |
| SLC44A4 | 6 | 16 | 1.39E-03 | 4.47E-02 |

\* Variants with internal study MAC $\leq 3$ were excluded from the analysis.

#### GoT2D Study

We performed single variant and gene-based tests with GoT2D deep exome sequencing data. Given that the ESP dataset contains few Finish samples (see Method), we focused on non-Finish GoT2D cohorts in this analysis, which included 650 cases and 646 controls. The gene-based rare-variant test results are summarized through QQ plots in Figure 3 (top panel); the iECAT-O QQ plot was fairly well calibrated. In contrast, iECAT-O_Noadj_ resulted in a significantly inflated QQ-plot. Even after the external control samples were ancestry matched (a total of 2568 external control samples; Web Figure 8), the QQ-plot by iECAT-O_Noadj_ remained significantly inflated, which suggests that technical batch effects were a major source of the type I error inflation (Figure 3, bottom panel). In contrast, iECAT-O had a well-calibrated QQ plot. Figure S9 compares p-values calculated with all 4300 ESP external control samples and 2568 ancestry-matched external control samples. These two p-values were largely consistent, indicating that iECAT-O analyses with and without ancestry-based matching were largely similar in this dataset.

We also performed single variant tests (Web Figure 10) for low frequency and rare variants with MAF < 0.05. As expected, QQ plots of iECAT were well calibrated, whereas QQ plots of iECAT_Noadj_ were greatly inflated. When we compared allele frequencies between the internal control samples and the matched external control samples, approximately 13% and 23% variants had p-values < 10% and 20%, respectively. These 3% inflation may indicate that approximately 3% of variants were subject to technical batch effects. To investigate whether the batch effects vary by minor allele counts (MAC), we obtained a distribution of the shrinkage weight *τ* by MAC (Web Figure 11). The box plots were similar across all four MAC bins, indicating that there was no clear pattern in batch effects by MAC.

Web Table 4 presents the top 5 genes by iECAT-O. Although GoT2D analysis did not identify statistically significant T2D-associated genes, this analysis provided a suggestive association and demonstrated that the proposed approach can control for type I errors.

## Discussion

In this paper, we proposed rare variant tests that increase power by integrating external control samples. By estimating bias using an Empirical Bayesian approach, the proposed iECAT method provides an effective way to adjust for possible batch effects. The type I error simulation and GoT2D data analysis revealed that iECAT can control for type I errors in the presence of technical batch effects. The power simulation and AMD data analysis revealed that iECAT can improve power compared with the exclusive use of internal controls. The method is implemented in the iECAT R package (available on the author’s website).

One of the important features of iECAT is that it only needs allele counts from external studies. By utilizing allele count information readily available in variant servers, such as Exome Variant Server (ESP) and ExAC browser (Lek et al., 2016), our method can greatly facilitate the use of external control samples. This is a desirable property considering the difficulties in obtaining individual-level genotype data. One possible issue of using allele count information in a variant server is that we cannot filter out samples with different genetic background. In such case, iECAT could yield slightly inflated type I error rates. We believe that it will become less problematic in the future as variant servers are starting to provide allele counts for fine-scale ancestry.

There are several limitations in the proposed methods. First, currently the methods cannot adjust for covariates, such as age and gender. This issue will be addressed in future research. Second, the method assumes that HWE holds (Guedj, Nuel, & Prum, 2008). Since it is uncommon to observe rare allele homozygotes, the violation of HWE condition would have limited effects on the proposed tests. Third, we do not recommend using the method for singletons or doubletons because log odds ratio estimates can be unstable for these variants. In future work, we will extend the method to incorporate singletons and doubletons.

With the advances in sequencing technologies, the number of sequenced genomes is increasing rapidly. Our method provides a robust and effective approach to utilize these sequenced genomes in rare variant tests and will contribute to the understanding of the genetic architecture of complex diseases.

## Supplemental Data

Supplemental Data are comprised of web appendices, four tables and eleven figures.

## Acknowledgements

This work was supported by grants R00 HL113264 and R01 HG008773 (SL), and the Austrian Science Fund (FWF) grant J-3401 (CF). We would like to thank investigators of GoT2D project for access to the whole exome sequence data, and Dr. Goncalo Abecasis for access to the ESP data. The dataset used for the AMD analysis described in this manuscript were obtained from the NEI Study of Age-Related Macular Degeneration (NEI-AMD). Database found at https://dbgap.ncbi.nlm.nih.gov through dbGaP accession number [phs000684.v1.p1].

